# Permittivity-Based Microparticle Classification by the Integration of Impedance Cytometry and Microwave Resonators

**DOI:** 10.1101/2022.09.27.509785

**Authors:** Uzay Tefek, Burak Sari, Mehmet S. Hanay

## Abstract

Permittivity of microscopic particles can be used as a classification parameter for applications in materials and environmental sciences. However, directly measuring the permittivity of individual microparticles has proven to be challenging due to the convoluting effect of particle size on capacitive signals. To overcome this challenge, we built a sensing platform to independently obtain both the geometric and electric size of a particle, by combining impedance cytometry and microwave resonant sensing in a microfluidic chip. This way the microwave signal, which contains both permittivity and size effects, can be normalized by the size information provided by impedance cytometry to yield an intensive parameter that depends only on permittivity. The technique allowed us to differentiate between polystyrene and soda lime glass microparticles — below 22 microns in diameter— with more than 94% accuracy, despite their similar sizes and electrical characteristics. The technique offers a potential route for targeted applications such as environmental monitoring of microplastic pollution or quality control in pharmaceutical industry.

## Introduction

The characterization of microscale objects constitutes a key problem in biological, materials and environmental sciences. In biomedical applications, high-throughput techniques are needed to differentiate cell types^1^ or test antibiotic susceptibility.^2^ In materials science, the optimization of drug loading efficiency of micropolymers^3^ can increase the efficiency of medications and lower their cost of production. In environmental screening, the widespread usage and high mobility of industrial microparticles have recently raised concerns about their environmental impact^4^ especially within the context of microplastic pollution.^5^

For the material identification of microscale objects, several analytical techniques, such as Raman and Fourier Transform Infrared Spectroscopy (FTIR), are in use. Even though these techniques can provide accurate material characterization, they are regarded as expensive, non-portable, and time-consuming for microparticle analysis.^6,7^ On the other hand, there are high-throughput techniques which provide the geometric size of microparticles —but not their material characteristics— such as resistive pulse sensing^8^ (commonly used in Coulter counters and nanopore analyzers) and more generally impedance cytometry.^9^ By augmenting impedance cytometry with the capability of material characterization, the applications mentioned above can be streamlined significantly.

In impedance cytometry, an analyte particle (such as a cell or microparticle) is detected by the change in the ionic current when it passes through the proximity of the sensing electrodes. At lower frequencies (<1 MHz, typically), the response of a dielectric particle is similar to an ideal insulator: it partially blocks the path of ionic current between the electrodes. The blockage current is proportional to the volume of the particle, providing a means for sizing microparticles. At radio frequencies (typically 1 to 50 MHz in practice), the capacitive response of the dielectric particle starts to contribute to the impedance signal, opening a potential route for material characterization. Indeed, several approaches in recent years have utilized the combination of a low-frequency (∼1 MHz) and high-frequency (∼50 MHz) impedance measurement to differentiate non-biological microparticles (e.g., polystyrene microparticles) from biological cells.^2,10-15^ In most of these studies, however, the differentiation was facilitated by the high-water content of cells: since water has a very large dielectric constant (ε_r_=78 at 1 MHz), it significantly affects the impedance signals. As a result, the high-water content of biological particles —such as cells— renders them to be readily distinguishable from any other non-biological particles which do not contain water. Critically, this advantage ceases to exist in differentiating amongst particles which do not contain water, such as two inorganic microparticles. In this case, the impedance signals generated are very close to each other, as discussed in detail in the Theory section below. Thus, leaving aside the specific case of biological particles, the dielectric-based differentiation of different microparticles in water poses a formidable challenge in sensing technology.

While impedance cytometry has gained widespread acceptance as a technique for determining the geometric size of particles, another electronic technique has been under development for microparticle analysis: microwave resonant sensors.^16-32^ By operating at the microwave frequency range (GHz frequencies), these sensors can bypass the Debye shielding effects across material interfaces and directly probe the capacitance of microparticles.^33,34^ When a particle passes through the active sensing region, the capacitance of the microwave resonator is modified. The modulation in the capacitance can be detected in the phase/amplitude response of the microwave resonator. In this work, to overcome the challenge of differentiating non-biological microparticles in a microfluidic system, we combined the two electronic sensor technologies mentioned above on the same system: a low-frequency (∼500 kHz) impedance cytometry sensor^2,35-37^ and a high-frequency (∼5 GHz) microwave capacitance sensor (Fig. 1). The vast frequency difference enables the two sensors to provide two parameters complementary to each other. The impedance cytometry sensor detects the geometrical volume of the particle within the channel;^11^ whereas the microwave sensor^16-23,26,27,30^ yields capacitance which is a function of the geometrical size and the Clausius-Mossotti factor of the particle, a factor that depends on the particle’s electrical permittivity and provides a means for material differentiation based on permittivity differences.

**Fig. 1.**
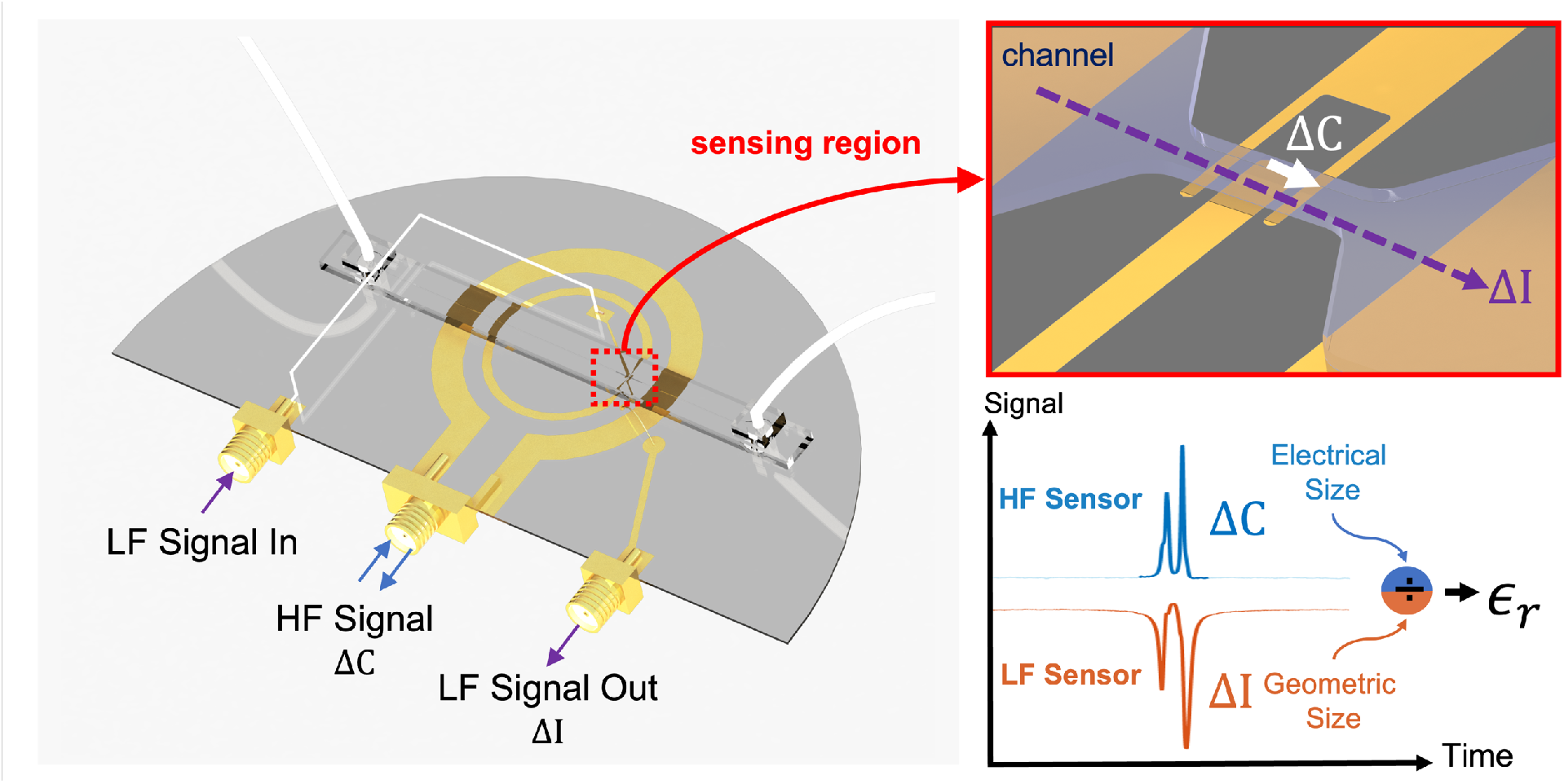
Sensing concept. On the microfluidic platform, there are two electronic sensors, a low-frequency (LF) sensor and a high-frequency (HF) microwave sensor. As a particle passes through the constriction on the microchannel collocated with the capacitive gap of the microwave sensor, it induces signals that encode its geometric size (current change in the LF sensor), and electrical size (capacitive change from the HF sensor). By combining electrical and geometric size, the permittivity of each particle can be deduced. In addition to the current and capacitance signals, the velocity of the particle can also be measured with this system which is used to factor out the vertical positional dependency of the microparticle in the channel.

In this paper, we introduce a novel, multiphysical sensor that provides a rapid, cost-effective, and highly accurate method for identifying and characterizing microscale particles, including glass and plastic. Our sensor employs a post-processing algorithm to eliminate errors caused by variations in position dependency, enabling it to detect and classify microplastics based on their electrical permittivity. This information can be used to gain insight into the sources and pathways of microplastics in the environment. We evaluated our sensor using two common types of non-biological microparticles: polystyrene (a common microplastic) and soda lime glass (a common household glass), both approximately 20 micrometers in diameter. With these analytes, our hybrid sensor attained signal-to-noise ratio of more than 200 in the microwave measurements, and 120 in low-frequency current measurements. With these high-resolution levels, our experimental results indicate that the ratio of Clausius-Mossotti Factors of the two microparticles is calculated to be 1.08, which is very close to the theoretical value of 1.10. Despite the low intrinsic contrast of the microparticles, our microfluidic sensor achieved 94.9% accuracy in identifying particle materials. Overall, the proposed sensor has the potential to significantly advance research in material and environmental science, enabling rapid identification and characterization of microscale particles.

### Positional Dependency of Electronic Signals

The precision of the measurements in such coplanar electrode designs —both for impedance cytometry and microwave sensors— is affected by the vertical position of particles within the analyzed volume, leading to a considerable degree of measurement inaccuracy.^12^ Microparticles of the same size but positioned at different heights within a channel exhibit different signal amplitudes due to the non-uniform electric field generated by coplanar electrodes. Therefore, relying solely on the raw magnitude of the electronic signal is inadequate for accurately sizing and quantifying microparticle samples.

Positional-dependent responsivity can be addressed by different approaches. One approach is to guide the particles to specific altitudes within the channel by the use of inertial focusing or microscale hydrodynamic effects such as sheath flow.^38-42^ However, these techniques may require high flow rates or add complexity to the sensor operation. A more recent approach to compensate for the positional effects involves the use of electrically floating electrodes to generate a prescribed variation in the electric field based along the vertical direction. In one such work, a differential measurement setup that utilized seven electrodes was developed.^37^ The outermost four electrodes are used for position calibration, while the middle three electrodes enable high signal-to-noise ratio current measurement. Alternatively, a five-electrode configuration was employed for position calibration and particle size measurements were performed concurrently by fitting the signal to a template.^35^ In both of the studies, a parameter called prominence (P) was used to calculate the height of each particle and calibrate its response. This parameter is observed^35^ to correlate with the velocity of the particle in the microfluidic channel, which is also verified in our system (Supplementary Information Fig. 2). Thus, we measured the velocity of the particles in our system by detecting the particle at two sensing locations, this way we could determine the height of the particle in the channel and calibrate the response accordingly. The calibration of particle height in both low-frequency and microwave sensors enables the separation of different particles.

**Fig. 2.**
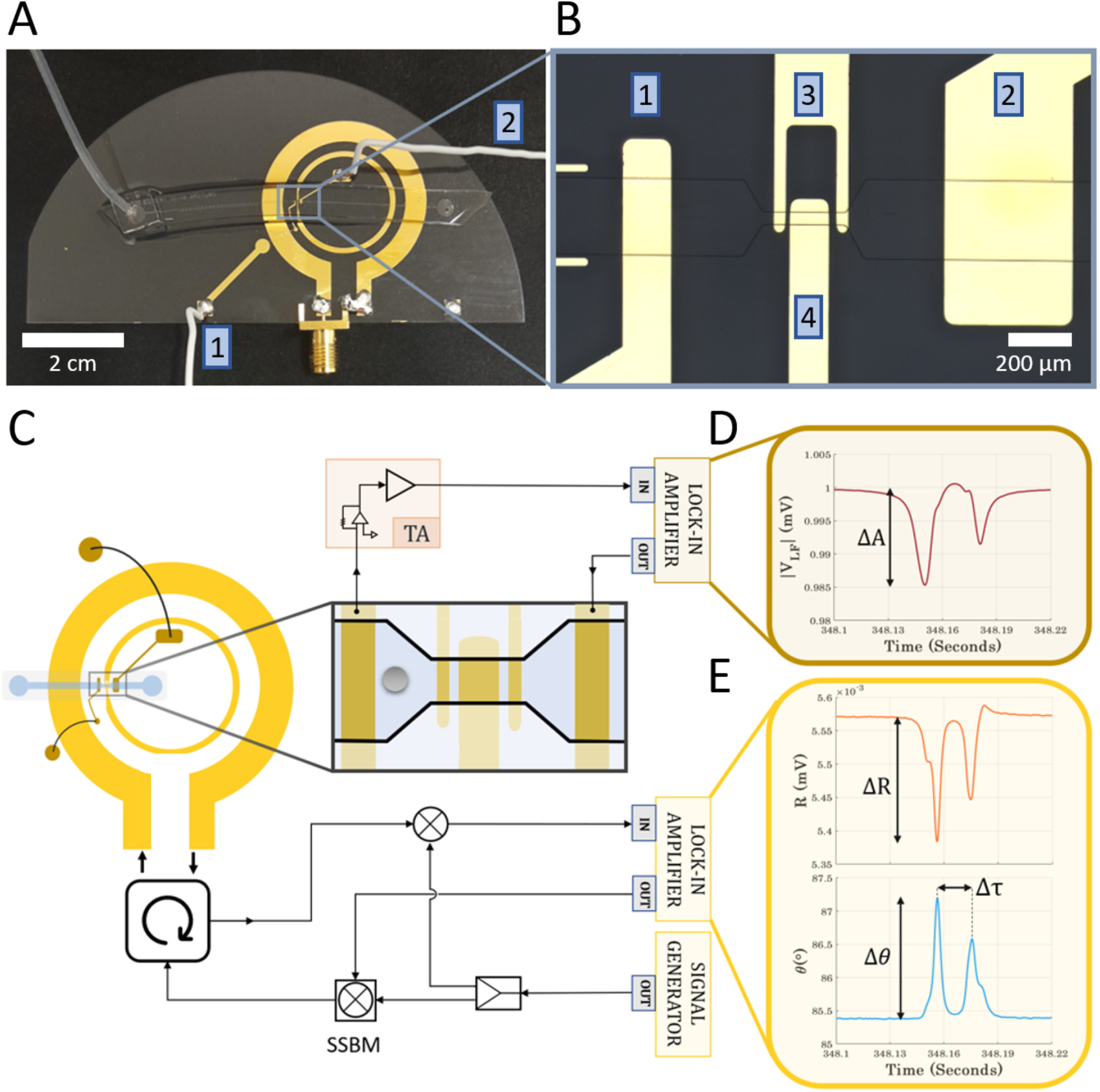
(**A**) A photograph of the sensor depicts: the microfluidic channel, the split ring resonator (SRR) for microwave sensor terminated by an SMA connector, gold tracks for the low-frequency sensor which are wire bonded or soldered to the pads to pass over the rings of the SRR. (**B**) A micrograph of the sensing region. The electrodes 1 and 2 are connected to the corresponding tracks in part (A). The low-frequency excitation is provided from the 2^nd^ electrode and collected from the 1st electrode. The electrodes 3 and 4 are for high-frequency (microwave) and particle velocity measurements. (**C**) Simplified schematic of the circuit, SSBM: single side-band modulator, TA: Transimpedance Analyzer. (**D**) Low-frequency current signal of a passing particle forms peaks when entering and exiting the microfluidic constriction. The geometric size of the particles is calculated from the initial peak. (**E**) Microwave signals induced by a passing particle. The initial changes in amplitude (ΔR) and phase (Δ*θ*) values are used to calculate the capacitance change induced by the particle.

### Theory

The operation principles of both sensor technologies integrated here have been verified independently in earlier studies. For the low-frequency sensor, the analyte particle induces an impedance change across a pair of electrodes in a conductive solution at 500 kHz. At low frequencies, a dielectric particle acts as an insulating object, partially blocking the ionic current between the electrodes. The amount of current blockade depends on the geometric size of the particle in the channel, rather than its material properties such as permittivity.^8,9,43^ Therefore, by measuring the change in low-frequency current passing through electrodes, one can obtain the geometric volume (*V*_*p*_) of the particle: Δ*I* ∝ *V*_*p*_. The second measurement is the high-frequency (microwave) capacitance measurement Δ*C* which generates a peak signal^16-19^ which depends both on the geometric size of the particle and its Clausius-Mossotti factor (K_CM_). K_CM_ is defined as:

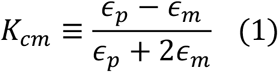

where ε_p_ stands for the permittivity of the particle, and ε_m_ that of the medium. By taking the ratio of the microwave signal (Δ*C*) to the low-frequency signal (Δ*I*), we obtain a quantity that depends on the Clausius-Mossotti factor of a particle, but not its volume.

Once K_CM_ is obtained, the permittivity of the particle can be back calculated by using the known permittivity value of the medium. In the determination of the microwave capacitance change Δ*C*, the large dielectric constant of water smooths out the differences in the response of different particles. As an example, consider the two classes of materials investigated in this work: polystyrene (ε∼ 2.55) and soda lime glass (ε∼ 7.2). While the ratio of their permittivity values is ∼3, this apparently large ratio does not translate into the contrast between impedance signals. Rather, the measured signals mostly reflect the dielectric properties of the equivalent volume of water that each particle replaces in the sensing region, as dictated by the large permittivity of the medium (ε_m_) in the Clausius-Mossotti formula (Equation 1). When the medium is water, the Clausius-Mossotti factors turn out to be –0.47 for polystyrene and –0.43 for soda lime glass: two close values differing by only 10% of each other. To put this in perspective, it is instructive to conduct a similar comparison for the differentiation of cells from microplastics. The relative permittivity values of cells range between 40-60;^44,45^ taking ε_r,cell_ = 50 for concreteness, the Clausius-Mossotti value turns out to be –0.1 for a cell. This value is almost five times smaller than the K_CM_ of polystyrene. In other words, a polystyrene particle induces approximately five times larger signal compared to a single cell of identical size; making it easy for differentiation from a cell. By contrast, the signal of polystyrene is only slightly larger (10%) than that of a micro glass particle of identical size. Such small differences between the Clausius-Mossotti factors create a challenge for the use of impedance cytometry in environmental screening to differentiate *anhydrous particles from other types of anhydrous particles* (*e*.*g*., plastics *versus* glass particles). To overcome this challenge, we integrated two high-resolution electronic sensors, which operate at frequencies separated by four orders of magnitude (500 kHz *vs. 5* GHz) on the same microfluidics platform.

### Experimental Procedure

Standard microfabrication and soft lithography techniques were used to fabricate a glass/PDMS microfluidic system with a gold layer serving as the electrodes for both the low-frequency and microwave sensors. The fabricated device (Fig. 2A) contains a microfluidic channel for analyte transportation, two gold tracks as the electrical ports of the low-frequency sensor, and one SMA connector for microwave sensing. The two concentric circles visible in the picture form the microwave resonator, called a split ring resonator (SRR) explained in more detail below ^23,46,47^. The sensing part of the device consists of two regions as shown in Fig. 2B: A) Two electrodes (labelled as 1 and 2) on either side for low-frequency measurements to obtain the geometric size of particles; B) Two microwave electrodes (labeled as 3 and 4) serving the dual purpose of enabling both microwave capacitance and particle velocity measurements.

Optical microscopy was used to observe the passage of particles through the sensing region concurrently with electronic measurements. For the analyte transportation, we used a PDMS microchannel which was sealed by the sensor chip. The microchannel was pressurized by Fluigent MFCS-EZ pressure control system and the analyte particles were passed through a constriction where the high frequency electrodes were placed. The purpose of the constriction was to enhance the resolution of the current measurement signal and to reduce the passage of multiple particles simultaneously from the sensing region. The dimensions of the constriction were 40 µm in width and 45 µm in height. The flow rates through the constriction ranged from 8 µL/min to 15 µL/min. Typically, the transit time of particles between two microwave gaps falls within the range of 10 to 20 milliseconds.

The low-frequency part of the sensor operates as a classic Coulter counter. The 2^nd^ electrode (Fig. 2B) is used to provide an AC excitation voltage to create a current through 1^st^ electrode. This current is carried by the ions in the solution, and a non-conducting particle passing through the channel blocks the ionic motion, resulting in a drop in the current proportional to the particle’s volume. To minimize the Debye screening effect without creating a coupling with high-frequency signal, the exposed surface area of the 1^st^ and 2^nd^ electrodes is maximized. The occurrence of two peaks in the signal is attributed to the presence of underlying gold electrodes in the microfluidic constrictions where the particle-induced current blockage is effectively circumvented by the transfer of electric current via the gold pads, rather than through the liquid medium. Consequently, the effect of particles on the low-frequency signal is neutralized while they pass through the microwave electrodes (Fig. 3B).

**Fig. 3.**
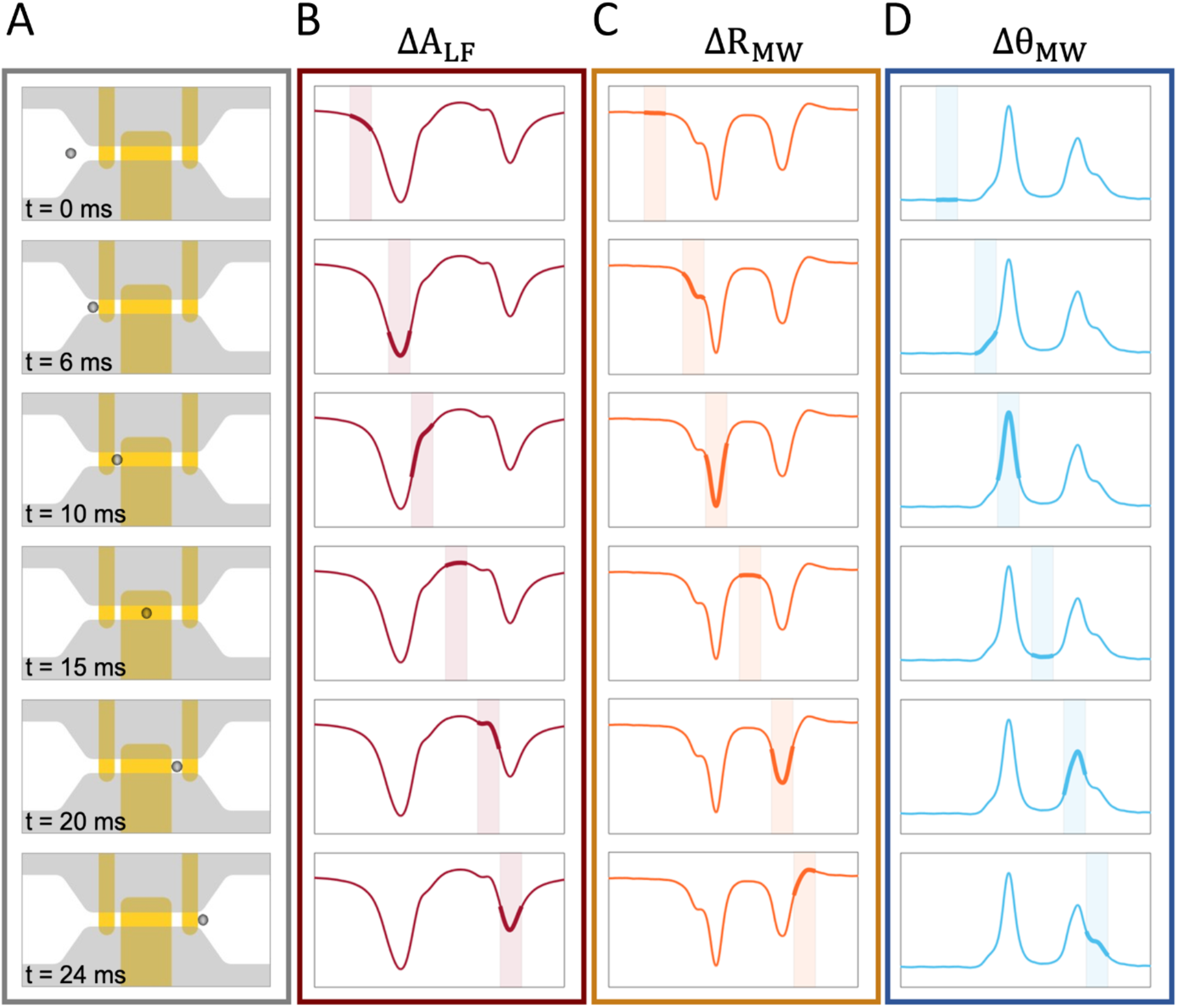
Progression of the particle through the sensing region and generation of signals. (**A**) Shows particle’s position schematically in a 24 ms time interval. (**B**) Shows the time domain raw data of a singular event’s low-frequency signal, with the corresponding section of each time window highlighted. The initial manifestation of the particle’s effect is observed at t = 0, coinciding with the narrowing section of the microchannel width. This effect reaches its maximum magnitude when the particle is fully inside the constricted zone at t = 6 ms and is subsequently neutralized while passing over the gold electrodes. The constriction length is designed to create a brief exit section without any gold layer beneath it, resulting in a smaller second peak in the low-frequency measurement. (**C**-**D**) Shows the microwave (MW) amplitude (ΔR) and phase (Δ*θ*) measurements. Two distinct peaks occurs while the particle is passing through the two gaps between the electrodes of the microwave resonator at t = 10 and t = 20 ms.

For the size measurement by the low-frequency sensor, the 2^nd^ electrode is driven by a 1 V peak-to-peak signal at 500 kHz from a lock-in amplifier (Zurich Instruments, HF2LI). The resulting current is collected from the 1^st^ electrode and converted into voltage by a transimpedance amplifier (Zurich Instruments, HF2TA), and read out by a lock-in amplifier (Zurich Instruments, HF2LI). The final signal, denoted as Δ*A*, is expressed in volts. For electrical size measurements at microwave frequencies, we used a split ring resonator^46-48^ (SRR) which consists of two concentric rings. The microwave signal was fed through the outer ring which inductively excites the inner one. Since there was a split along the inner ring, a standing-wave mode shape emerges and creates a high intensity electric field in the split region.^46-48^

To ensure the accuracy of measurements, it is essential to have only one particle present in the constricted section at any given time, as signals from multiple particles interfere with one another. To this end, we implemented a data analysis technique that detects and disregards overlapping particle passages. To facilitate this technique, we used two distinct gaps (15 and 25 µm wide) located within the inner circle of the split ring resonator. The utilization of two different gap sizes is critical in distinguishing whether a secondary peak in time is caused by the passage of the same particle through the second gap or another particle from the first gap. The narrower gap generates a more concentrated electric field, thereby inducing a greater change in capacitance which was subsequently used for the electrical size calculation of the particles under investigation. Conversely, the wider gap induces a smaller capacitance change. In the end, each particle creates two distinct changes on both amplitude and phase values; a relatively large signal induced in the narrow gap and a smaller signal caused by passage through the wider gap. By measuring the time delay between the signals from the two gaps, the velocity of the particles is calculated and used to calibrate the signals to obtain the electrical and geometric size accurately.

After verifying the resonance characteristics of the SRR with a vector network analyzer (VNA), we switched to our custom circuitry (Fig. 2C) and set the frequency of the signal generator to the resonance frequency (∼5.4 GHz). The circuit consists of two lock-in amplifiers to employ single side band modulation (SSBM) and a signal generator as signal sources. We fed the SRR at its resonance frequency with 500 mV peak-to-peak output. The time constant of the lock-in amplifiers was set to 501 µs and the sampling rate was 13.39k/sec to fully detect rapidly passing particles. Since the maximum operational frequency of the lock-in amplifiers were below the resonance frequency, we built a custom Single Side Band (SSB) heterodyne circuitry ^16,17^. The reference signal outputs of the lock-in amplifiers were up-converted before entering the resonator and down-converted back before digitally reading the amplitude and phase of the signal. A detailed explanation of the circuitry can be found in the Supplementary Fig. S3 and a detailed explanation of the data analysis can be found in the Materials and Methods section.

The particle diameter was determined from the low-frequency signal (Δ*A*), while the velocity and height information of each particle were extracted through the delay between two consecutive peaks on the phase data. For the geometrical diameter *D*, the signal depends on the maximum signal value: 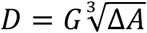. Here *G* is a calibration constant of the circuitry, which can be determined with the help of a calibration particle whose precise diameter is known. In our circuitry, this constant was empirically found to be 88.9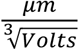.

After the low-frequency sensor, we analyzed the signal from the microwave sensor. To obtain the capacitance change, we monitor the amplitude (R) and phase (*θ*) response of the microwave resonator to calculate the out-of-phase component (*i*.*e*., *Y* quadrature, with *Y* ≡ *R sinθ*) of the reflected voltage from the microwave resonator. Any fractional change in the capacitance of the microwave resonator is reflected in the out-of-phase component (Supplementary text 1):

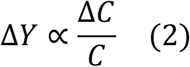

Here, Δ*Y* denotes the change in the out-of-phase component of the reflected voltage, and *C* denotes the total capacitance of the microwave resonator. The apparent values (before height compensation) of Δ*R* and Δ*θ* the particles were extracted for each event. The velocity was then used to compensate for the effect of the trajectory height of each particle by:

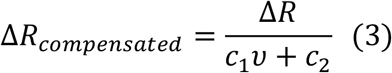

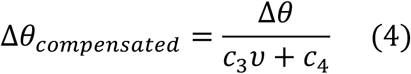

The determination protocol for constants (i.e., *c*_1_ *c*_2,_ *c*_3,_ *c*_4_ are explained in detail in Supplementary Fig. 5. By using these two parameters, the capacitance change induced by a particle can be calculated by the relationship:

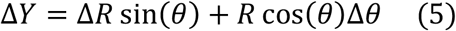

where *R* and *θ* correspond to baseline value of raw signals. This way, the two-sensor system provides both geometric diameter and the height-compensated electrical area values which can be used for material classification. Since the same constants are used in the compensation protocols in equations (4) and (5), the separability of the different materials is not affected by this procedure.

## Results

With this experimental procedure, we compared the measurements from two types of microparticles: polystyrene and soda lime glass. We used the large geometric span of the soda lime glass particles (Cospheric, SLGMS-2.5, 15-22 *µ*m size distribution) to infer the scaling law between the impedance cytometry and microwave measurements in a loglog plot (Fig. 4A). The slope of the loglog plot was 0.646 ± 0.014 (*i*.*e*., approximately 2/3) which indicated that the microwave signal scaled with the 2^nd^ power of diameter, whereas the impedance cytometry signal with the 3^rd^ power, to give a ratio of 2/3 which is close to the observed slope. Hence, to obtain the geometrical diameter, we calculated the cube root of the impedance cytometry signal, and for the electrical diameter, we calculated the square root of the microwave signal (Equ. 6) and plotted them in Fig. 4B. In this plot, we have both the soda lime glass particles in blue, as well as the polystyrene particles in orange, which have a much narrow diameter distribution (20 ± 0.3 µm). Based on the observed scaling trend, the slope on this chart is proportional to the square root of K_CM_ value for each particle.

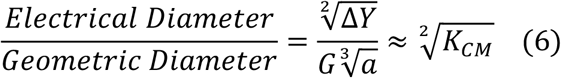

**Fig. 4.**
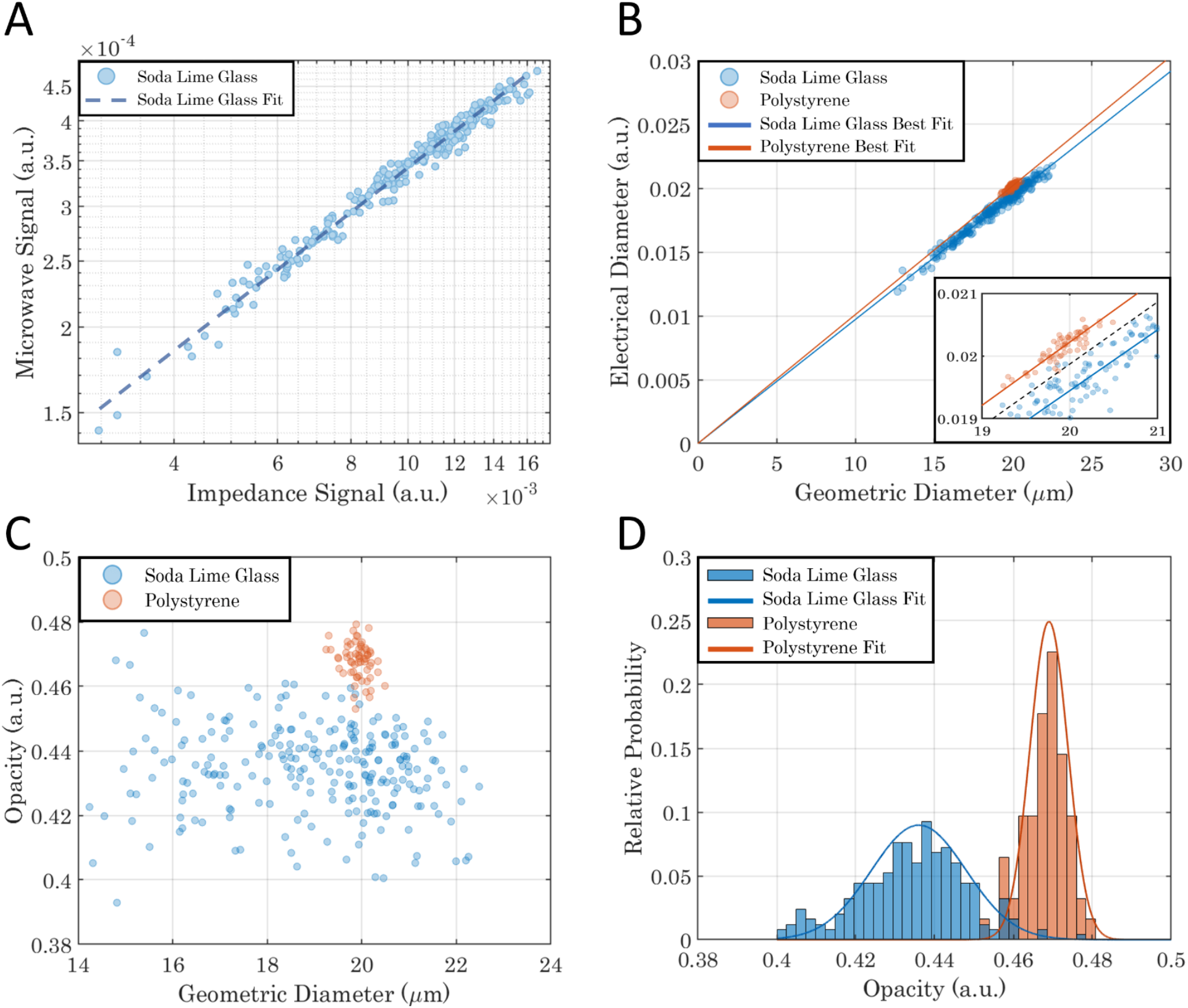
(**A**) The velocity-calibrated signals from impedance cytometry (x-axis) and microwave sensors (y-axis) of soda lime glass particles on logarithmic scale. (**B**) Separation of polystyrene and soda lime glass using geometric and electrical diameters. (**C**) Separation by the opacity of polystyrene and soda lime glass (**D**) Histogram of opacity showing material classification along with separating Gaussian fits.

The separation of the slopes of the two particle types is evident in Fig. 4B. As expected, polystyrene particles have higher |*K*_*CM*_| and this fact is reflected in Fig. 4B as they have a sharper slope compared to soda lime glass (|*K*_*CM*_| factors of polystyrene and soda lime glass are 0.47 and 0.43, respectively). The black line in Fig. 4B inset shows the separation region between the polystyrene and soda lime glass. Within this binary separation protocol, the technique achieves correct separation with 94.9% accuracy overall.

Fig. 4C shows the opacity of the particles at microwave frequencies which is a parameter used to determine the amount of electrical field penetration^2^ a particle allows for. It appears from the results that polystyrene beads are opaquer to electrical field penetration. This is indeed also expected because of the permittivity contrast with water. Whenever there is an interface in fluid where ions are present, these ions block the field penetration (i.e., Debye Screening). At microwave screening where Debye screening no longer affects signal magnitude, the opacity value is expected to overlap with the Clausius Mossotti factor. By calibrating the opacity with the known Clausius Mossotti factor of polystyrene, they resulted in 0.47 (σ =5.5 10^−3^) for polystyrene and 0.43 (σ =1.4 10^−2^) for soda lime glass. These values were found to be consistent with the expected ratio. The histogram in Fig. 4D indicates that this opacity difference can be used to classify different dielectric materials. We can also use the statistics of the polystyrene group only (which has a narrow size distribution), to determine the resolution of the technique. In both geometric and electrical diameter measurements, the ratio of the standard deviation to the mean value turns out to be 1.2%. With this attained high resolution in individual parameters, their ratio —which is proportional to the opacity factor— also yields a narrow distribution which enables the differentiation of the two materials which have very similar values (0.47 *vs*. 0.43). Material specific misclassification rates are detailed in Table 1.

**Table 1.** Statistics of Analyzed and Classified Dielectric Microparticles.

|  | Number of Particles | Number of<br>Misclassifications | Successful Separation<br>(%) |
| --- | --- | --- | --- |
| Polystyrene | 62 | 2 | 96.8 |
| Soda Lime Glass | 249 | 14 | 94.4 |
| Total | 311 | 16 | 94.9 |

Apart from the challenging differentiation of dielectric microparticles, we also verify that the integrated approach presented here can readily differentiate a non-biological particle from a biological cell; or a dielectric microparticle from a metallic microparticle as shown in Supplementary Fig. 6. This way, different classes of materials, in addition to different dielectric materials, can be quickly and reliably distinguished with this approach.

## Conclusion

In conclusion, we have shown that two different classes of dielectric microparticles that have very similar electronic characteristics can be differentiated from each other based on their permittivity. To achieve the differentiation capability, we integrated two different electronic sensors that operate at frequencies separated by four orders of magnitude: a 500 kHz sensor for geometric sizing and a microwave (∼ 5 GHz) sensor for electrical sizing. By combining the two signals, we obtained an intrinsic material property for each microparticle and reached an identification accuracy of 94.9% in the binary classification of polystyrene and glass microparticles.

While this technique holds great promise for materials and environmental science applications, it is important to note some limitations of our study. Our current approach is limited to the classification of materials with distinct dielectric permittivity values. A sensor with higher resolution can enable the differentiation of microparticles amongst a larger variety of material classes, which will be of immense benefit for the materials and environmental sciences and their applications. Furthermore, the use of a lower permittivity medium rather than water may result in signals with higher contrast for microparticles of different material content. By using a medium with lower permittivity, the difference in K_CM_ values can be maximized, leading to more distinguishable permittivity values for different microparticle materials. Further research can explore the potential benefits of using a lower permittivity medium in the identification of different microparticle materials. Nevertheless, the potential implications of our findings are significant. By enabling the differentiation of microparticles among a larger variety of material classes in water, this approach could be applied to various fields, including the identification of pollutants in the environment and quality control in manufacturing.

## Materials and Methods

### Fabrication of the Sensor

The sensor device was fabricated on one half of a 100 mm x 0.5 mm (diameter x thickness) double-side polished wafer. The substrate was chosen to be dry fused quartz, a non-conductive and low loss-tangent material. Sensor geometry was patterned by spin coating 2 µm thick AZ – 5214 photoresist and photolithography steps. After the photoresist development in AZ – 400K for 30 seconds, 5 nm of chromium and 150 nm of gold were deposited by the thermal evaporation process. Chromium was used as a binding agent between quartz and gold layers. After an overnight acetone lift-off process, the sensor pattern was completed. For the microchannel fabrication, SU-8 mold was used to create channel patterns on a PDMS block. After punching the fluidic inlet and outlet through the PDMS block, the bottom surface of the channel and the gold deposited wafer was treated with oxygen plasma to activate their surfaces. Within ten minutes after the activation, the microfluidic channel and the wafer were aligned and bounded under an optic microscope

### Data Analysis

In our experimental setup, we recorded changes in both the current level at low-frequency current and the reflected voltage wave at microwave frequency. The first step in analyzing the collected data was to discern single events, which posed a challenge due to the presence of peaks of varying sizes and those that were closely spaced. As such, the raw data was unsuitable for standard MATLAB peak finding algorithms. To address this issue, we utilized a two-step convolution process on the microwave phase (*θ*_*MW*_) signal. The outcome was the transformation of single event signals into a Gaussian-shaped signal, as demonstrated in Supplementary Fig. 4. Initially, the phase signal was convolved with the sign function. The length of the function was chosen to fully cover a single particle event signal. In the case of multi particle passage, the convolution operations interfered with each other and resulted in asymmetric shapes which ended up with multiple peaks after the Gaussian smoothing process. After calculating the exact time windows of the events and filtering out multi peak shapes, the standard peak finder algorithm was able to identify all the signal values accurately. In our study, we opted for a sign function length of 15 ms to avoid excessive interference in the low-frequency signals. This means that only if two events occurred consecutively with a delay of more than 15 ms, the convolution operation yielded a single peak.

### Medium Preparation

To increase the magnitude of the current, and maintain compatibility for particle/cell characterizations, we used 0.2% Tween 20 (Sigma-Aldrich, P1379-500ML) added Phosphate-buffered saline (Biowest, L0615-500) as the medium in the experiments which has a conductivity of 9 mS/cm. Polystyrene particles were received in DI water: a 3 µL volume of this solution was mixed with a 5 mL volume of the Tween 20 / PBS solutions. Soda lime glass particles were received as a solid powder and dissolved in the same Tween 20 / PBS solution.

### Calibration and Reliability

The two types of particles had similar diameters and optical appearances, making it difficult to identify them by optical microscopy at high flow rates used in the measurements. For this reason, particles were passed through the sensor system sequentially to have an independent verification of the identity of each particle. The microwave resonance curves were compared between the runs to ensure that device sensitivity was not changed. The tests were repeated on a second device with identical dimensions, and the separation between the two class of microparticles was verified with similar accuracy level.

## Data Availability

All the data supporting the findings of this study are available within the paper and its Supplementary Information files or from the corresponding author upon request.

## Acknowledgements

The authors thank Hashim Alhmoud, Ceren Alatas, Berk Kucukoglu, Hadi S. Pisheh and Arda Secme for useful discussions. This project has received funding from the European Research Council (ERC) under the European Union’s Horizon 2020 research and innovation programme (grant agreement No 758769).

