## Supplementary Text and Figures for "Permittivity-Based Microparticle Classification by the Integration of Impedance Cytometry and Microwave Resonators"

### Dielectric Classification at the Single Microparticle Level by Microwave Sensing Integrated with Impedance Cytometry

**Supplementary Text 1: Mathematical Modelling**

The microwave aspect of the sensor is modeled as a series RLC circuit where the particle passage in the microchannel can be modeled as a capacitance change of ΔC (Fig. S1). In such circuits, the response of the circuit can be expressed as a transfer function in frequency domain. For a single port design like our SRR, the transfer function corresponds to the ratio of reflected wave voltage $V^-$ to the incident wave voltage $V^+$. This ratio is also defined as $S_{11}$ parameter. This parameter is a function of line impedance of the measurement circuitry, and input impedance of the device $Z_{in}$.

$$H(\omega) = \frac{V^-}{V^+} = S_{11} = \frac{Z_{in} - Z_0}{Z_{in} + Z_0}$$

For and RLC circuit the transfer function can be written as,

$$H(\omega) = \frac{(R - Z_0) + j\left(\omega L - \frac{1}{\omega(C + \Delta C)}\right)}{(R + Z_0) + j(\omega L - \frac{1}{\omega(C + \Delta C)})}$$

Since the sensor is always driven at its resonance frequency, the impedance of the sensor has to be purely real. Therefore, at resonance.

$$\omega_{res} L - \frac{1}{w_{res} C} = 0$$

Also, if use Taylor Expansion to the capacitance term to the first degree;

$$\frac{1}{\omega_{res}(C + \Delta C)} = \frac{1}{\omega_{res} C} - \frac{\Delta C}{\omega_{res} C^2}$$

Then, the transfer function at resonance frequency can be written as;

$$H(\omega_{res}) = \frac{(R - Z_0) + j\frac{1}{\omega_{res}}\left(\frac{\Delta C}{C^2}\right)}{(R + Z_0) + j\frac{1}{\omega_{res}}\left(\frac{\Delta C}{C^2}\right)}$$

If the expression is multiplied with its complex conjugate and it's real and imaginary parts are separated.

$$Re\{H(\omega_{Res})\} = \frac{R^2 - Z_0^2 + \frac{1}{\omega_{res}^2}\left(\frac{\Delta C}{C^2}\right)^2}{(R + Z_0)^2 + \frac{1}{\omega_{res}^2}\left(\frac{\Delta C}{C^2}\right)^2}$$

$$Im\{H(\omega_{Res})\} = \frac{\frac{2Z_0}{\omega_{res}}\left(\frac{\Delta C}{C^2}\right)}{(R + Z_0)^2 + \frac{1}{\omega_{res}^2}\left(\frac{\Delta C}{C^2}\right)^2}$$

The capacitance change is going to reflect itself to the first order in the out-of-phase component (i.e., the imaginary part of the transfer function), but the second order in the in-phase component (i.e., the real part of the transfer function). Furthermore, $\Delta C \ll C$ which means $\left(\frac{\Delta C}{C^2}\right)^2 \ll 1$, and the imaginary part can be simplified as;

$$Y = Im\{H(\omega_{Res}, \Delta C)\} = \frac{\frac{2Z_0}{\omega_{res}C}}{(R+Z_0)^2}\left(\frac{\Delta C}{C}\right)$$

**Fig. S1.**

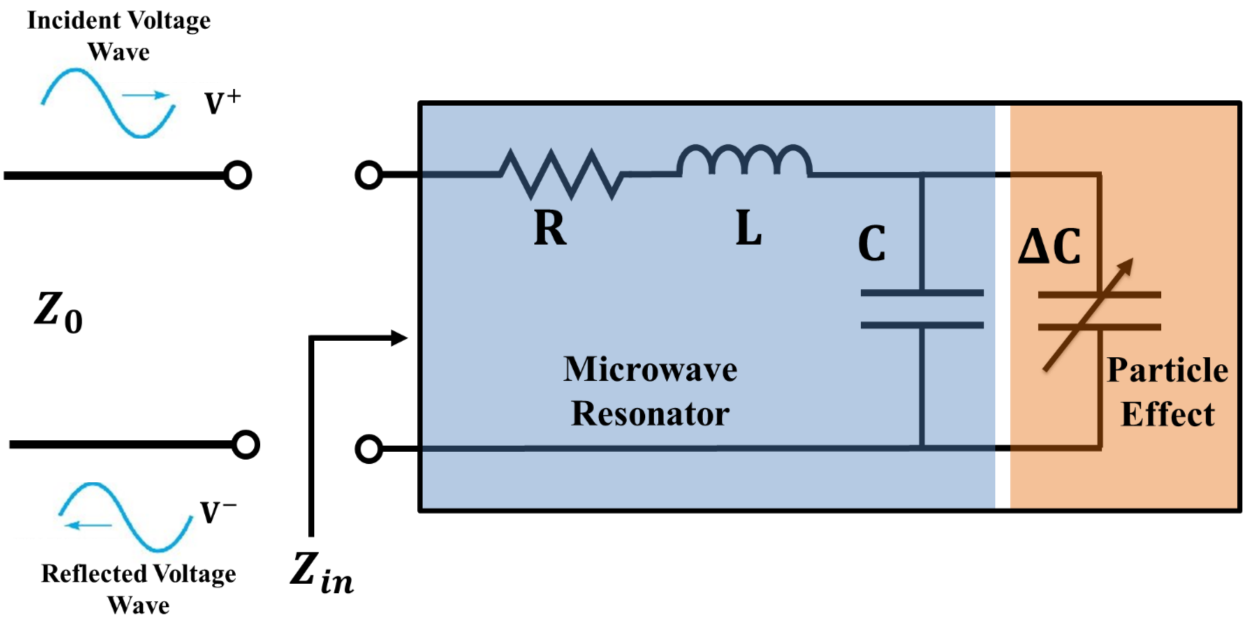


**Fig. 1S. Microwave Resonator Modelling:** Particle passage in the microchannel can be modeled as a capacitance change of ΔC to the RLC circuitry.

**Fig. S2.**

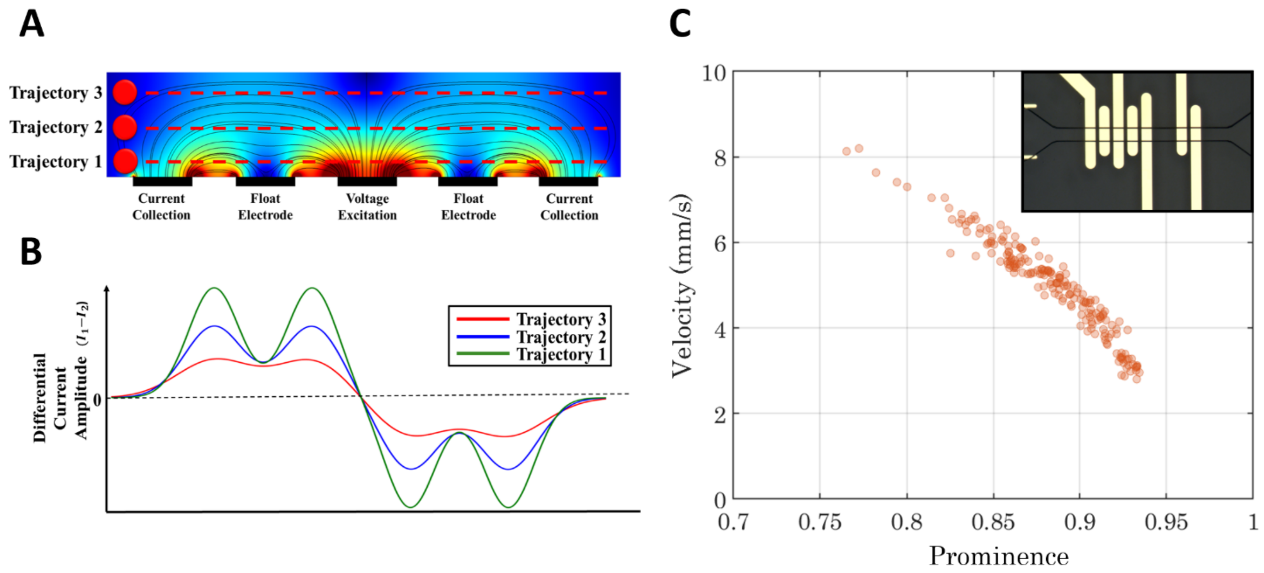


**Fig. S2. A set of experiment to point out the relation between velocity and height:** (**A**) Cross section of a five-electrode, differential measurement design as employed in.[35] Black rectangles are the electrodes and thin black lines correspond to equipotential points. Particles flow through the channel from various heights. In this schematic, three height values are used. Each trajectory experiences different magnitude of electric field along its path which enables us to calculate the height. (**B**) A particle passing closer to the electrodes induces a large variation in the current along its trajectory, whereas the current waveform is smoother for an identical particle passing near the top of the channel. The waveform is encoded as a height dependent parameter called 'Prominence' which corresponds to the ratio between the large and small peaks in the signal waveform. (**C**) Demonstration of prominence and velocity relation for 20 $\mu m$ polystyrene particles. The five electrodes on the left are for prominence calculation, and the two on the right are for microwave sensing.

**Fig. S3.**

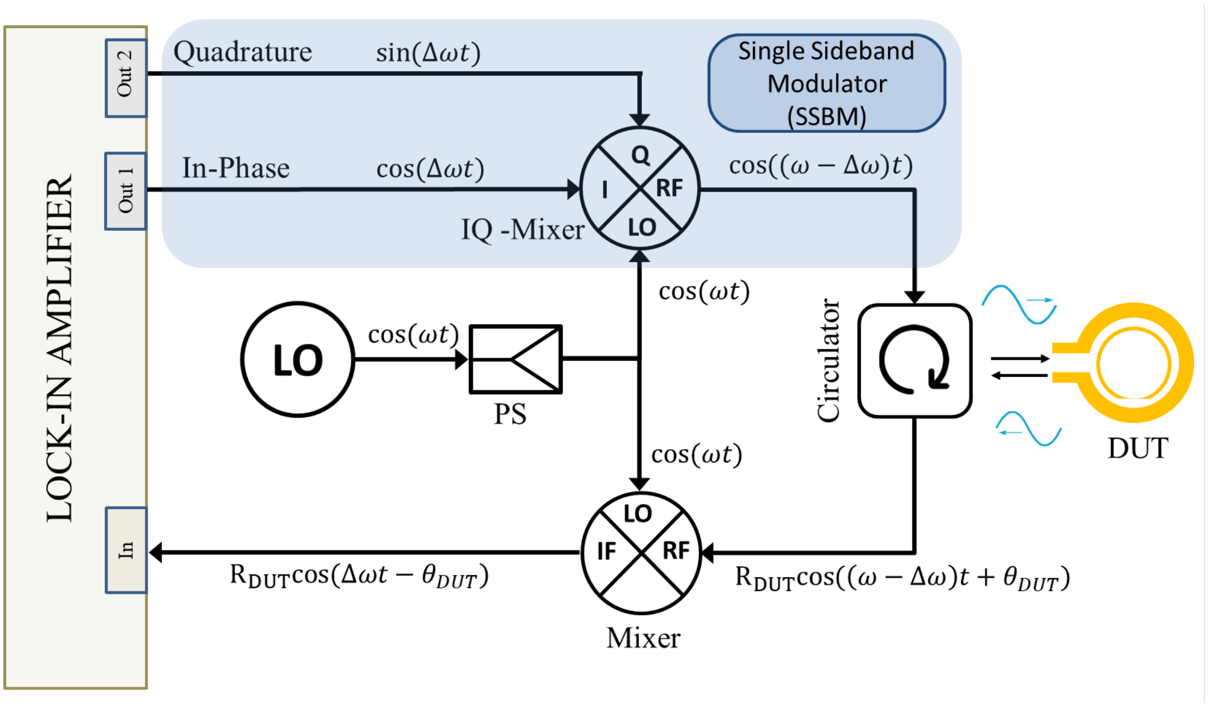


**Fig. S3. Main custom circuitry diagram, showing up-mixing and signal reading in detail:** The custom-built measurement circuitry consists of mainly a single sideband modulator and a down conversion stage where a mixer is employed. Lock-in amplifier (Model Number: ZI MFLI) is used to create a low frequency (~300 kHz) signal. In order to excite the microwave resonator at the resonance frequency, we use a signal generator (i.e., local oscillator, LO) to provide the high frequency (~5 GHz) signal. To inject the low frequency signal onto microwave, signal an IQ mixer (Model Number: MMIQ-0218LPC) is fed with two outputs of the lock-in amplifier which are separated by 90° phase difference, yet at the same frequency. As a result of this mixing with the LO signal, we get a signal at the frequency which is the frequency difference between signal generator (R&S SMB100A) and lock-in amplifier. This signal is then fed to microwave resonator via a circulator. Since our resonator is a single port microwave device a circulator is employed to direct incident and reflected voltage waves from the resonator. The reflected voltage wave is then directed to a mixer where it is downconverted back to kHz frequencies. Here, the same signal that is fed to IQ mixer is used as the local oscillator source for the mixer. After down conversion the low frequency signal is analyzed by lock-in amplifier where the amplitude and phase are digitally sent to the computer.

**Fig. S4.**

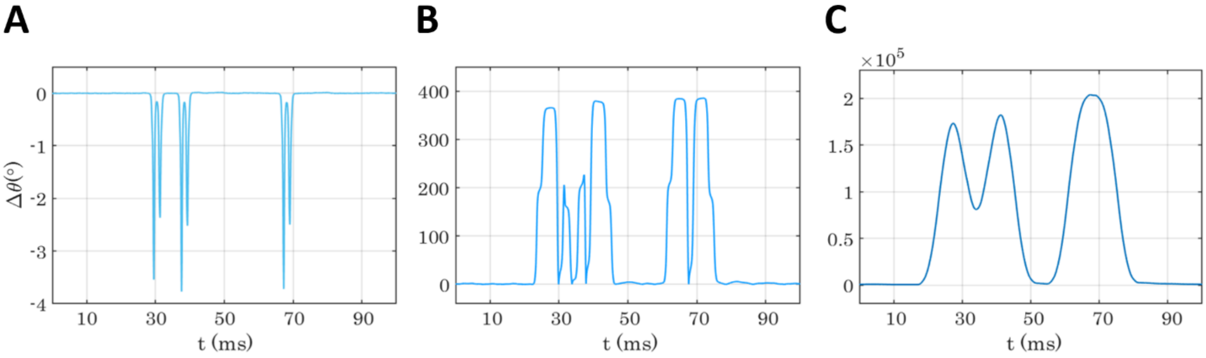


**Fig. S4. Two step convolution process on Phase signal to detect single events:** (**A**) A time domain, phase data of three events. The third event is isolated enough but the first two are too close to each other for a reliable characterization. (**B**) The outcome of the first convolution operation. While the first two events produced two peak pairs with different shapes, the third produced a symmetric pair. (**C**) The result of the second convolution (Gaussian smoothing) operation. The first two events merged into a multi-peak shape easily detectable with a standard MATLAB peak finder algorithm.

**Fig. S5.**

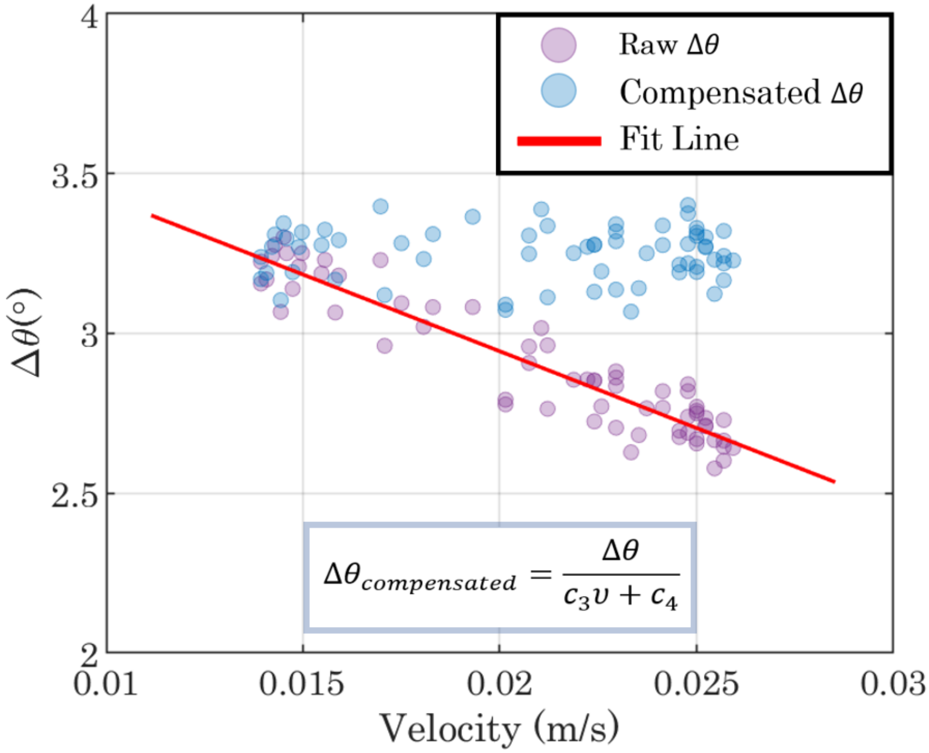


**Fig. S5. Compensation Operation:** In high frequency sensing, each particle is measured in two distinct locations, which results in two signal peaks on both phase and amplitude parameters as it can be seen in the main text Figure 3. This method allows determination of both electrical volume and travelling velocity of a particle in the microchannel. Since particle trajectory height is dependent on its velocity, the compensation operation is based on velocity. In order to have a precise compensation, we use 20 $\mu m$ polystyrene particles as the calibration particles. We start the compensation operation by first calculating the velocity of each polystyrene event by measuring the time delay between the event peaks. To find parameters used in main text (i.e $c_1$(-0.022), $c_2$(0.00015) for ΔR; $c_3$(-538), $c_4$(3.9) for $\Delta\theta$), we fit a linear function to polystyrene data. These parameters are reported in the table below. Since we use normalized diameter in fitting, same linear fit can be used for compensation of all particles of various diameters.

**Fig. S6.**

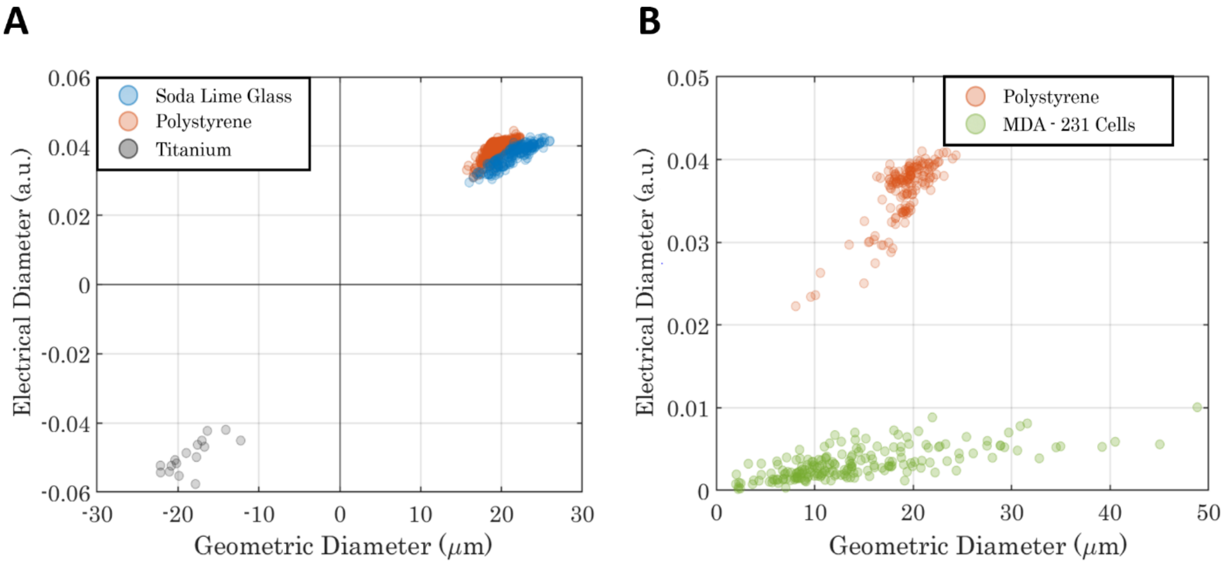


**Fig. S6. Differentiation of dielectric microparticles from metallic microparticles and non-biological particles from biological cells:** (**A**) Differentiation of different material classes can be readily achieved compared to the differentiation amongst dielectric materials. The signal of Titanium microparticles can be quickly differentiated owing to the opposite sign they generate in both sensors. Since metal particles facilitate, rather than impede the electric current, their blockage diameter is interpreted with a negative sign. (**B**) The comparison of polystyrene microparticles and single cells obtained with a device with simplified architecture (without any compensation element). In this sensor, only one high frequency measurement (instead of two) were made.
